# NMR-based analysis of nucleotide π-stacking in a crowded environment: Implications for prebiotic reactions

**DOI:** 10.1101/655423

**Authors:** Niraja V. Bapat, Harshad Paithankar, Jeetender Chugh, Sudha Rajamani

## Abstract

The inherent heterogeneity of the prebiotic milieu is often overlooked when studying nonenzymatic reactions. However, it is important to note that the prebiotic soup of a putative ‘RNA World’ would have been replete with a plethora of molecules resulting from complex chemical syntheses, as well as exogeneous delivery. The presence of such background molecules could lead to pertinent phenomenon such as molecular crowding, which can potentially affect how a reaction would advent in a crowded milieu. In the current study, we have analyzed the effect of crowding on the stacking ability of the RNA monomers, using Nuclear Magnetic Resonance (NMR) spectroscopy. Our findings corroborate that the purine monomers possess better stacking efficiency than pyrimidine based monomers. Significantly, this competence is further enhanced in the presence of a crowding agent. Interestingly, this enhanced stacking could result in higher sequestration of the purine monomers, putting their ready availability for relevant nonenzymatic polymerization and replication reactions into question. Taken together, this study demonstrates the need for systematic biophysical characterization of molecular crowding in the context of prebiotically pertinent processes. Unravelling such phenomena is essential to gather a real understanding of how the transition from abiotic to biotic, would have happened during the origin of life.

## Introduction

Chemical origins of life posit that the prebiotic soup would have facilitated important chemical reactions, which are thought to have led to the origin of life on early Earth. This composite solution would have been replete with different kinds of molecules, whose reactivity and interactions would have impinged on the processes that led to the emergence of the first cell-like entities. The heterogenous nature of the soup could have been, partly, due to the highly diverse range of products that would have resulted from various prebiotically relevant syntheses. It is well-known that the outcome of most of such reactions is a mixture of products. For example, the formose synthesis reaction that results in a mixture of 4 to 6 carbon containing sugar molecules, never exclusively yields only one kind of sugar (Zweckmair et al., 2014). Along with D-ribose that is found in RNA molecules, this reaction also yields a plethora of other sugars such as erythrose, xylose, arabinose, glucose, mannose etc. Another example is the Fischer-Tropsch-Type (FTT) synthesis pathway, which is considered as a prebiotically viable means of synthesizing longer carbon chain compounds from carbon monoxide and hydrogen. Significantly, FTT reactions are also known to yield a a complex mixture of alkanols, alkanoic acids, alkenes, and alkanes, containing anywhere from 2 to over 30 carbon moieties (McCollom et al., 1999). Similarly, only a fraction of the resultant product from Or?’s synthesis, is constituted of the canonical adenine nucleobase (Orgel, 2004). On the other hand, the meteorites that would have fallen on the prebiotic Earth, during the ‘Late Heavy Bombardment period’, would have been comprised of a mixture of compounds. It is pertinent to highlight that the analysis of Murchison meteoritic samples has confirmed the presence of a plethora of sugar-related compounds and organic molecules (Cooper et al., 2001; Schmitt-Kopplin et al., 2010).

All such examples clearly indicate that the early Earth would have been rife with different kinds of molecules that would have been present concomitantly in the prebiotic soup. Therefore, any prebiotically pertinent reaction could not have taken place in isolation and under buffered conditions, unless completely secluded by some relevant mechanism. The existence of various types of molecular entities (co-solutes or background molecules) would have resulted in a complex prebiotic milieu that, consequently, would have facilitated phenomenon inherent to such scenarios, like molecular crowding, which has been shown to affect, both, the kinetics and equilibrium of several biochemical reactions (Ellis, 2001). Furthermore, macromolecular crowding has been shown to impact the protein folding, stability, and rates of the enzymatic reactions that they catalyze (Minton, 2001; Zhou et al., 2008). Significantly, it has also been demonstrated to influence the reactions involving nucleic acids (Nakano et al., 2017). In the presence of water-soluble crowding agents, the thermal stability of DNA duplexes (ranging from 8-30 mer) is known to be dependent on, both, the DNA length, and the size of the co-solute (Nakano et al., 2004). A few studies have also delineated the effect of molecular crowding on prebiotically relevant molecules like ribozymes. For example, catalytic RNAs are known to get stabilized under crowded conditions (Kilburn et al., 2013; Desai et al., 2014), and show enhanced activity in the presence of co-solutes (Nakano et al., 2009; Kilburn et al., 2013; Strulson et al., 2013; Desai et al., 2014).

In addition to the direct interactions, there is another conceivable way of explaining the effect that crowding agents could have on macromolecule-based reactions. Crowding could potentially alter molecular diffusion that is precipitated by the presence of bulky co-solute polymers. Crowding agents are known to reduce the diffusion coefficient (D) of, both, small and large molecules (Ellis, 2001). Diffusion of the monomers would have played an important role in polymer formation during the origin of life. Interestingly, restricting monomer diffusion using certain methods, such as surface adsorption on clay mineral (Ferris 2006), entrapment of monomers in dried lipid films (Toppozini et al., 2013; Himbert et al., 2016), and crowding of monomers in the liquid channels within eutectic ice phases (Kanavarioti et al., 2001; Monnard et al., 2003) etc., has been demonstrated to enhance the yields of nonenzymatic polymerization reactions.

In this study, we have analyzed the changes in the diffusion, and thereby the stacking properties, of RNA monomers in aqueous solution, upon addition of a molecular crowding agent. To our knowledge, this study is possibly the first of its kind wherein RNA monomer diffusion has been evaluated, especially in the prebiotic context. Nuclear Magnetic Resonance (NMR) spectroscopy was used for this purpose as it is a non-invasive technique and does not require any tagging of the molecules in question. Polyethylene glycol (PEG), which is a bulky water-soluble inert polymer, and has been used to simulate crowded conditions in several biochemical studies (Zhou et al., 2008), was used as a molecular crowding agent in these experiments. Our observations indicate that the purine monomers stack to a greater extent than pyrimidine monomers, under crowded conditions. This could affect their availability for participating in prebiotically relevant reactions, such as nonenzymatic polymerization and replication of nucleic acids. In a particularly relevant study, it was reported that PEG and DLPC lipid vesicles, when used as co-solutes, reduced the rate of purine-based addition reactions in template-directed copying of RNA. This further resulted in an overall reduction in the fidelity of the copying of information in this RNA-based system (Bapat & Rajamani, 2015). Essentially, this study highlighted the possibility of how such processes could have fundamentally impacted events that would have facilitated the transition from chemistry to biology.

## Materials and Methods

### Chemicals

The disodium salts of all four 5’-nucleoside monophosphates (5’-NMPs), viz. adenosine 5’-monophosphate (5’-AMP), guanosine 5’-monophosphate (5’-GMP), uridine 5’-monophosphate (5’-UMP), and cytidine 5’-monophosphate (5’-CMP) (all purity ≥ 98%), were purchased from Sigma Aldrich (Bangalore, India) and used without any further purification. Analytical grade polyethylene glycol (PEG) 8000 and deuterated water (D_2_O) were also purchased from Sigma Aldrich.

## Methods

### Sample preparation

The nucleotide stocks were prepared in nanopure water and the concentrations were estimated using UV spectroscopy (UV-1800 UV/Vis spectrophotometer, Shimadzu Corp., Japan). A 50% w/v stock solution was prepared for PEG 8000 by dissolving the required amount of powder in nanopure water. Three different concentrations of the nucleotides viz. 10, 50, and 100 mM, were used to record the DOSY NMR data. For T_1_ relaxation time measurements, the data was recorded for nucleotide concentrations of 10, 40, and 100 mM. The final concentration of PEG 8000 in the analyzed samples was maintained at 18%. 300 μl of 1.1 times concentrated samples was first prepared and 10% D_2_O was subsequently added to these samples for field locking before recording the NMR data.

### NMR data acquisition

All the DOSY-NMR and T_1_ relaxation experiments were recorded on Acsend™ Bruker 600 MHz NMR spectrometer, furnished with quadruple (^1^H/^13^C/^15^N/^31^P) resonance Cryoprobe equipped with X, Y, Z- gradient; and dual receiver operating. DOSY experiments were recorded for the nucleotides in the presence and absence of PEG 8000. For each sample, the diffusion time and the gradient length was optimized to get 5-10% residual signal at 95% of the maximum gradient strength (42.58 G/cm). Sixteen data points were recorded with strength of the gradient ranging between 2-95%. The diffusion times (Δ), and the durations for which the gradient pulse (δ) was applied for all the analyzed samples, are listed in Table S1 in the supplementary data section.

For ^13^C-T_1_ relaxation time measurement, ^13^C signals were excited and detected on the attached protons via polarization transfer. The ^13^C data was recorded by providing six inversion recovery delays (5 delay points and a repeat point for error estimation) in the range of 100 to 800 ms. The recovery delays were chosen so as to get 70% reduction in the signal intensity at the longest delay. For all the samples (i.e. control samples as well as those containing 18% PEG 8000), the T_1_ relaxation data was recorded at two temperatures viz. 10 °C (cold) and 25 °C (hot).

### Data analysis

The collected DOSY data was processed using SimFit algorithm as explained in standard Bruker DOSY data processing manual (Kerssebaum, R. DOSY and Diffusion by NMR. In *User Guide for XWinNMR 3.5*, Version 1.0; Bruker Buospin GmbH: Rheinstetten, Germany, 2002). The intensities extracted by SimFit algorithm were fit using a two-parameter mono-exponential equation (described below) in OriginPro 8.5.0 (OriginLab, Northamptan, MA, USA) to get the diffusion coefficients for the solute molecule in the solvent that has been used in the study:

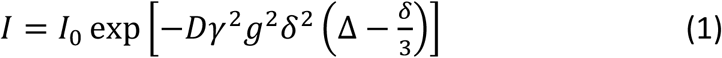

where, *I* is the observed intensity, *I*_0_ is the reference or un-attenuated intensity, *D* the diffusion coefficient, *γ* the gyromagnetic magnetic ratio of the observed nucleus, *g* the gradient strength, *δ* the length of the gradient pulse, and Δ the diffusion time.

The ^13^C-T_1_ relaxation data was processed and extracted as separate 1D corresponding to each recovery delay time, *t*. Peak picking was done in topspin3.2 (Bruker Biospin, GmbH: Rheinstetten, Germany). The intensity data was then fitted using Mathematica v5.2 (Wolfram Research, Inc., Champaign, IL, USA) script (Spyracopoulos, L. J.Biomol. NMR 2006, 36:215-224) to the following mono-exponential decay function, to get the longitudinal relaxation time constant (T_1_):

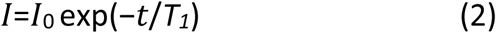

The error in the T_1_ time was obtained as fit error the to the above equation using both Monte Carlo simulations and repeat relaxation delay point from the Mathematica script.

## Results

### Translational diffusion slows down with increase in nucleotide concentration

To begin with, the DOSY NMR data was recorded for the nucleotide samples in the absence of PEG. The diffusion coefficients (D) obtained for these samples are listed in Table S2. Concentration dependent consistent decrease in D was observed for all four nucleotides, suggesting slowing down of the translational diffusion in the solution (Fig 1). However, the DOSY data recorded for nucleotide samples in the presence of PEG did not show any clear trend. This could arise since diffusion processes are governed primarily by the high viscosity of the solution in the presence of PEG. Hence, ^13^C-T_1_ relaxation experiment was subsequently used as a tool to compare the relative sizes of the molecules, in the presence and absence of PEG.

**Figure 1:**
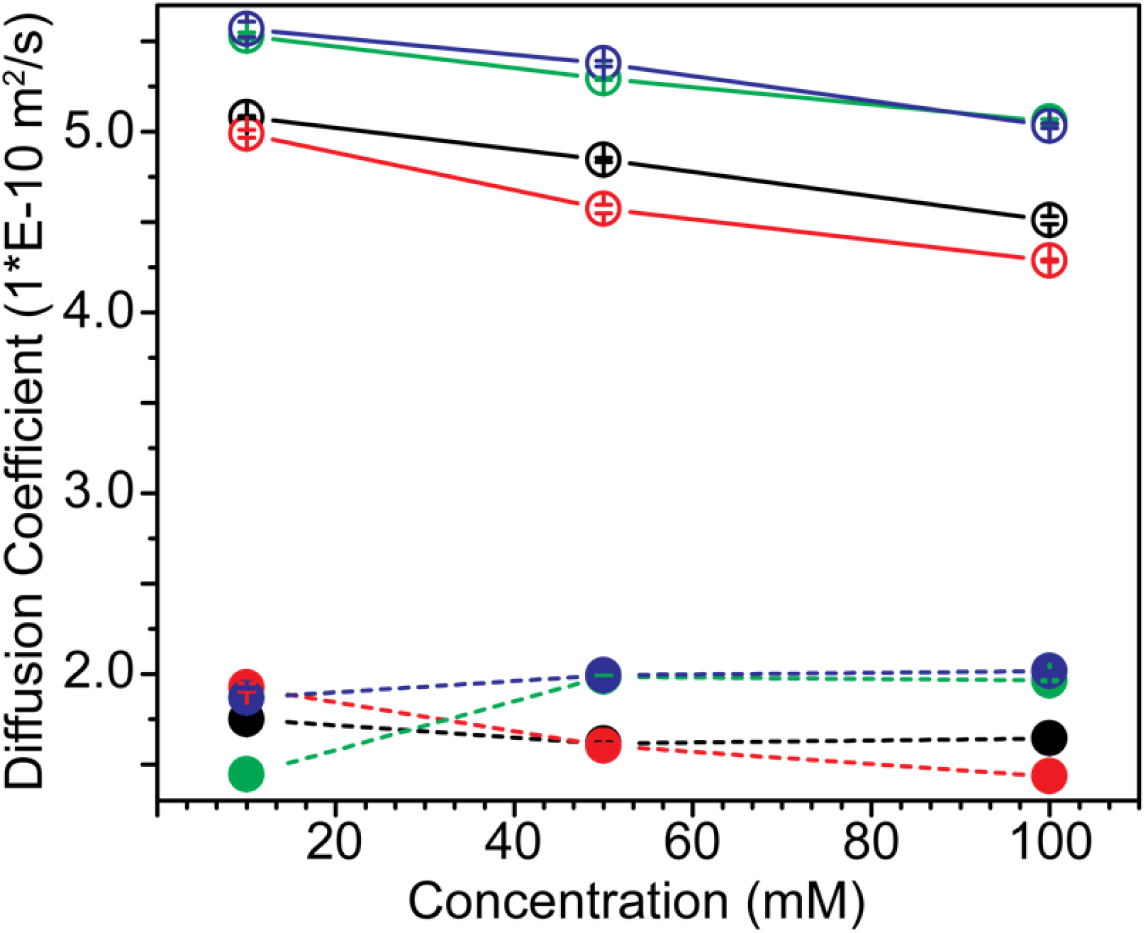
DOSY NMR measurements for nucleotide diffusion coefficients. The diffusion coefficients were measured at different concentrations, in the absence (indicated by empty circles connected by a solid line), and in the presence of PEG (indicated by filled circles, connected by a dashed line), for all four nucleotides viz. 5’-AMP (indicated in black), 5’-GMP (indicated in red), 5’-CMP (indicated in green) and 5’-UMP (indicated in blue). The X-axis indicates three different concentrations at which the data was recorded. Errors in the diffusion coefficients is shown by the bars and are of the order of 1%.

### Increase in the nucleotide concentration slows down rotational correlation time

The ^13^C-T_1_ relaxation data was recorded for all four 5’-NMPs at three different concentrations and at two temperatures viz. 10°C and 25°C. The theoretical T_1_ relaxation time follows a curve (defined by the equation below) passing through a minimum as the molecular size increases. Fig 2 shows ^13^C-T_1_ relaxation time plotted against the rotational correlation time of the molecule which is a determinant of the molecular size. The T_1_ values are obtained through simulations using following equations:

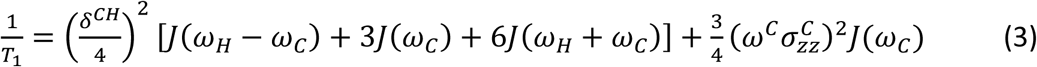

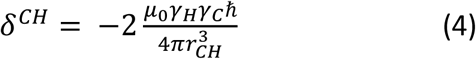

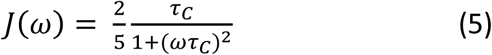

Where, *δ*^*CH*^ = Anisotropy of the dipolar coupling between C and H;

μ0 = Vacuum permittivity = 4π*10^-7^ N.A^-2^

*γ*_*C*_ and *γ*_*H*_ = Gyromagnetic ratio of ^13^C and ^1^H (6.728*10^7^ T^-1^.s^-1^ and 2.675*10^8^ T^-1^.s^-1^, respectively)

ℏ = reduced plank constant (1.05457*10^-34^ J.s)

*r*_*CH*_ = Distance between two nuclei C and H (1.1 Å)

*J(ω)* = Spectral density at frequency ω;

*ω*_*c*_ and *ω*_*H*_ = Resonance frequency of ^13^C and ^1^H;

*σ*_*zz*_ = Chemical shielding tensor; and

τ_C_ = Rotational correlation time of the molecule.

For the molecules with size characterized by left of the T_1_ minimum, T_1_ time increases with an increase in sample temperature, while for macromolecules which have size greater than that characterized by T_1_ minimum, relaxation time decreases with an increase in temperature (Levitt, 2001).

**Figure 2:**
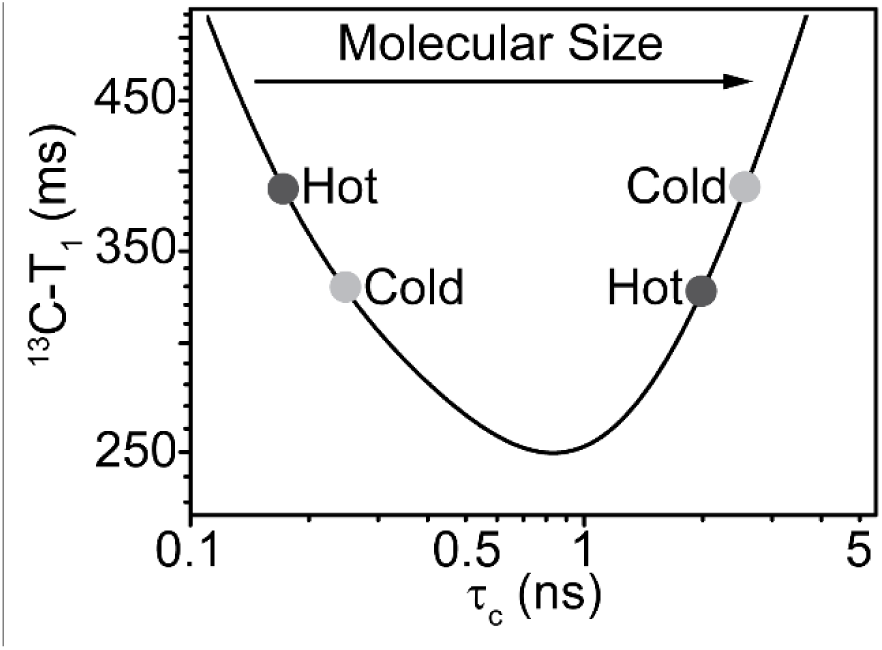
Log-log plot of ^13^C-T_1_ relaxation time as a function of molecular size (characterized by rotation correlation time(τ_c_)) shown as a temperature dependent quantity. Curve is simulated using equations 3, 4, and 5 to represent data measured at NMR spectrometer with magnetic field strength of 14.1 T, ω_H_ = 14.1*γ_H_, ω_C_ = 14.1*γ_C_, and chemical-shielding tensor (*σ*_*zz*_) = 0.00025 ppm.

The comparison of T_1_ relaxation times for all these samples showed an increase with increase in the temperature from 10 °C to 25 °C (Fig 3, Table 1). This suggested that the nucleotides behaved as small molecules (characterized by the left of the T_1_ minimum in Fig 2) under our analytical conditions. As the concentration of nucleotides was increased from 10 to 100 mM, a significant decrease in T_1_ relaxation time (ΔT_1_ ∼150 ms at 25 °C and ΔT_1_ ∼100 ms at 10 °C) was observed for purine monomers (5’-AMP and 5’-GMP). No such significant change was observed in the case of pyrimidine monomers (5’-UMP and 5’-CMP). The decrease in T_1_ for purine monomers with an increase in the nucleotide concentration suggested an overall increase in the rotational correlation time pointing towards increased stacking.

**Table 1:** T_1_ relaxation time data for different nucleotides in the absence and presence of 18% PEG 8000.

| Concentration of Nucleotide (in mM) | Without PEG |  | With PEG 8000 |  |
| --- | --- | --- | --- | --- |
| | $T_1$ (ms) at 10 °C | $T_1$ (ms) at 25 °C | $T_1$ (ms) at 10 °C | $T_1$ (ms) at 25 °C |
| **5'-AMP** |  |  |  |  |
| 10 | 468.48 ± 11.92 | 580.62 ± 14.94 | 271.54 ± 13.63 | 338.48 ± 10.87 |
| 40 | 361.69 ± 7.63 | 510.30 ± 8.26 | 287.24 ± 6.15 | 338.86 ± 3.57 |
| 100 | 326.10 ± 6.48 | 431.74 ± 4.39 | 271.16 ± 6.95 | 309.71 ± 12.16 |
| **5'-GMP** |  |  |  |  |
| 10 | 383.65 ± 22.77 | 469.11 ± 3.26 | 317.14 ± 15.44 | 280.72 ± 21.94 |
| 40 | 340.42 ± 4.87 | 429.74 ± 15.62 | 244.62 ± 7.48 | 323.08 ± 4.67 |
| 100 | 297.11 ± 2.90 | 369.39 ± 2.98 | 253.91 ± 6.78 | 315.45 ± 7.27 |
| **5'-CMP** |  |  |  |  |
| 10 | 391.23 ± 11.70 | 531.67 ± 5.75 | 289.84 ± 6.20 | 342.47 ± 18.37 |
| 40 | 380.12 ± 3.01 | 520.03 ± 6.81 | 284.39 ± 1.19 | 365.06 ± 6.87 |
| 100 | 361.04 ± 2.90 | 493.06 ± 4.55 | 271.92 ± 1.97 | 351.37 ± 4.49 |
| **5'-UMP** |  |  |  |  |
| 10 | 384.18 ± 5.39 | 547.65 ± 6.17 | 272.89 ± 6.57 | 385.46 ± 15.27 |
| 40 | 378.79 ± 2.81 | 522.71 ± 4.02 | 292.04 ± 6.19 | 369.22 ± 10.55 |
| 100 | 368.43 ± 2.04 | 506.53 ± 5.27 | 280.27 ± 1.94 | 370.59 ± 5.05 |

**Figure 3:**
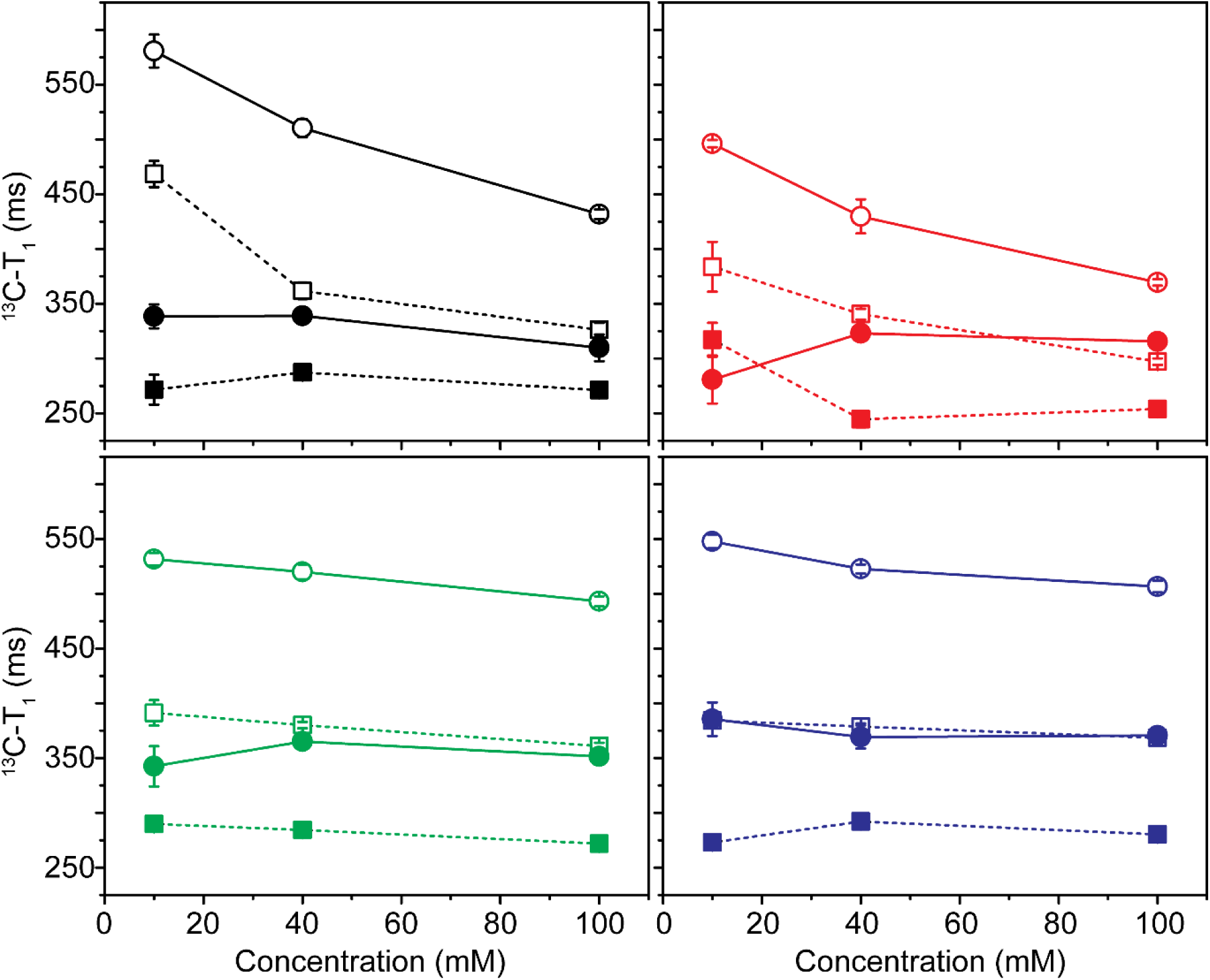
^13^C-T_1_ relaxation time data for different concentrations of nucleotides. Nucleotides are color coded as follows: 5’-AMP (Black); 5’-GMP (Red); 5’-UMP (Green); and 5’-CMP (Blue). The data was recorded for the nucleotides in the absence (empty shape), and in the presence of PEG (filled shape), at 25 °C (circle with solid line) and 10 °C (square with dashed line), respectively.

### Molecular crowding slows down rotational correlation time

The recorded T_1_ values were, in general, lower in the presence of PEG as compared to those obtained in its absence, due to the increase in the viscosity of the solution. In presence of PEG 8000, the T_1_ relaxation time showed a similar increase in (as in absence of PEG) from cold to hot measurement condition, implying the overall molecular size is characterized by left of the T_1_ minimum. This suggested that the nucleotides behaved as small molecules even in the presence of PEG. However, upon increasing the nucleotide concentration from 10 mM to 100 mM, T_1_ was found to decrease for purine monomers (except for 10 mM 5’-GMP) (Fig 3, Table 1), while it was invariant for pyrimidine monomers. This suggested an increase in the size of the nucleotide cluster/pseudo-oligomers in the presence of PEG.

## Discussion

The heterogeneous nature of the prebiotic milieu would have resulted in interesting phenomenon such as molecular crowding and related events. The presence of molecular crowding agents is known to affect the diffusion coefficient, D, of the solutes. If the D is reduced, molecules would take relatively longer to travel the same distance, thus resulting in reduced chances of encountering neighboring molecules, or their interacting partners. Hence, if the reaction is diffusion-limited, its rate would decrease in the presence of co-solute polymers (Ellis, 2001). It has been previously reported that the presence of PEG and lipid vesicles affect the rate and fidelity of template-directed nonenzymatic RNA copying reactions (Bapat and Rajamani, 2015). In this one of a kind study, the results underlined the importance of accounting for the presence of background molecules in prebiotically relevant reactions. This is especially pertinent as their presence could have directly impacted the kinetics of nonenzymatic oligomerization and replication processes, affecting the origin and the composition of informational polymers.

In the present study, we have analyzed the effect of molecular crowding on the diffusion of prebiotically relevant molecules viz. RNA monomers. RNA has been argued to be the first biomolecule to have arisen during the origin of life, due to its dual ability of carrying the genetic information and performing the catalysis (RNA World hypothesis, Robertson & Joyce, 2010). NMR analysis was used to probe the effect of PEG on ribonucleotide stacking, and thereby its effect on nucleotide diffusion, under our analytical conditions. Along with being a non-invasive method, NMR also bypasses the need to tag the nucleotides in order to track their movements. This is hugely advantageous as the attachment of bulky fluorescent tags would increase the effective nucleotide size, thus very likely affecting their stacking properties, and thereby their diffusion.

To begin with, DOSY NMR was the method of choice for this analysis as one can get a direct estimation of diffusion coefficients for molecules, under different solution conditions. In the absence of PEG in the solution, the diffusion coefficients showed a decrease with the concurrent increase in the concentration for all the four canonical 5’-NMPs, suggesting an increase in size of the stack of monomers with increase in concentration. Furthermore, the diffusion coefficient observed in the presence of PEG were lower in general; however, no concentration dependent decrease was observed. This might be due to the overall effect of PEG on the viscosity of the solution that levels the effect of nucleotide stacking on diffusion coefficients, such that no observable differences were detected in the diffusion coefficients, even at different concentrations. Importantly, spontaneous formation of columnar liquid crystals has been recently reported for, both, DNA and RNA monomers, at high concentrations and low temperature (Smith et al., 2018).

Since, the diffusion coefficients calculated from DOSY could not provide the differential size estimation for the various nucleotide concentrations studied in the presence of PEG, T_1_ relaxation time was subsequently recorded. T_1_ relaxation, which is also known as spin-lattice relaxation time, is a temperature-dependent function of the molecule, and hence can be used to estimate its size based on the rotational properties. In the absence of PEG, an increase in the T_1_ value was observed for the nucleotides, for a concurrent increase in the temperature from 10 to 25 °C. This suggested that the nucleotides acted as small molecules under this experimental set-up (Levitt, 2001), and lie on the left of the T_1_ minimum at all concentrations. However, a notable concentration dependent decrease in T_1_ was observed only for purine monomers at both the temperatures. This stems from better π-stacking properties exhibited by the purines, which might lead to the formation of pseudo-oligomers, ultimately resulting in larger sized molecule at higher concentrations. Pertinently, in a previous study, the diffusion coefficient was shown to be dependent on the number of nucleotides present in the ssRNA (Werner, 2010). Therefore, larger molecular size would result in a concurrent decrease in diffusion, such that the stacked purine monomers might be less available for the diffusion-limited reactions, as against what was observed for the pyrimidine monomers. Significantly, this effect could further get enhanced at higher concentrations of the nucleotides (Fig 3).

In the presence of PEG, the T_1_ relaxation time was observed to have decreased in general for all the nucleotides and at all concentrations. This could be primarily due to the increase in the viscosity of the solution in the presence of bulky PEG polymers. When the data was analyzed closely, it was noted that the overall decrease in the T_1_ values at 25 °C, compared to those observed at 10 °C, was lower for the purine nucleotides as compared to those obtained for pyrimidine nucleotides (Table 1). For example, the T_1_ for 100 mM 5’-AMP at 25 °C was found to be around 310 ms and that at 10 °C was ∼ 270 ms (the difference being that of ∼40 ms). On the other hand, the T_1_ for 100 mM 5’-UMP at 25 °C was found to be 370 ms and that at 10 °C was 280 ms (i.e. the difference of ∼90 ms). The reduction in the T_1_ value from higher temperature to the lower temperature would be less for a molecule/molecular cluster that is comparatively of larger size (Fig 2) (Levitt, 2001). Thus, this variation in the difference is suggestive of higher sized molecular clusters/pseudo-oligomers for purine monomers in the presence of PEG, than that for pyrimidine monomers. The molecular clusters/pseudo-oligomers formed by purines are concentration independent in the concentration range tested, as evidenced by no significant change in the increase in T_1_ times from 10 to 25 °C (Fig 3 and Table 1).Thus, PEG might facilitate higher degree of stacking for purine monomers, resulting in the sequestration of the monomers into these pseudo-oligomeric structures.

Another peculiar thing that was observed from the relaxation data recorded in the presence of PEG was the T_1_ values for 5’-GMP. For 10 mM of 5’-GMP, the recorded T_1_ relaxation time was higher at 10 °C than at 25 °C, in the presence of PEG (Table 1). This meant that in the presence of PEG, at lower concentration, 5’-GMP acted as a large molecule as estimated by the T_1_ relaxation data. However, at higher concentrations of 40 and 100 mM of 5’-GMP, the recorded T_1_ values in the presence of PEG were again found to be lower at 10 °C (Table 1). This indicated that at higher concentrations, the 5’-GMP cluster acted as small molecules. This is interesting because molecular crowding induced by PEG is known to promote formation of non-canonical G-quadruplex structures that result from the formation of non-Watson-Crick interactions between guanine (Miyoshi et al., 2006). We, therefore, suspected that in the presence of PEG, 5’-GMP could be forming G-quadruplex-like structures at higher concentrations, resulting in the compaction of the nucleotide clusters, and thus yielding a T_1_ relaxation time trend characteristic of a small molecule. To delineate this, ^1^H NMR analysis was carried out to check for the presence of G-quadruplex-like structures at higher concentrations of 5’-GMP in the presence of PEG. The formation of hydrogen bonds in G-quadruplexes results in the stabilization of exchangeable hydrogen, which can be observed as peak at ∼10-12 ppm on ^1^H NMR. The peak corresponding to the presence of G-quadruplex was observed at 40 mM 5’-GMP and was absent at 10 mM 5’-GMP (Fig S1). This observation supported the T_1_ data by confirming the formation of compact G-quadruplex like structures at higher GMP concentration, as compared to the looser stacks present at lower GMP concentrations.

Taken together, both the DOSY NMR and T_1_ relaxation time data indicate a higher propensity for stacking of the purine monomers, as against the pyrimidine monomers. Furthermore, the T_1_ data also indicated that this stacking tendency increased in the presence of PEG 8000. The effect of molecular crowding on π-stacking might result in the formation of pseudo-oligomers. This is especially prominent in the presence of PEG as opposed to plain aqueous solution conditions. Consequently, this would result in the sequestration of a higher percentage of purine monomers in such aggregates. These results explain, in part, the previously observed reduction in the rate of purine-based cognate nonenzymatic addition reactions, particularly in the presence of PEG (Bapat & Rajamani, 2015). Significantly, the observations from the aforementioned and current study, demonstrate the importance of accounting for prebiotic heterogeneity, and discerning its implications while characterizing pertinent nonenzymatic copying reactions. This is crucial as the presence of co-solutes could directly impinge on the propagation of genetic information during the origin of life on Earth. Overall, this study underscores how heterogeneous mixtures, and the emergent phenomena that they facilitate, could have direct consequences for characterizing pertinent prebiotic processes.

## Supporting information

NMR_NT stacking_Supplementary File

## Acknowledgments

The authors wish to acknowledge the High Field NMR facility at IISER-Pune (co-funded by DST-FIST and IISER Pune). JC acknowledges Department of Biotechnology, Govt. of India [BT/PR24185/BRB/10/1605/2017], and the Science and Engineering Research Board (SERB), Govt. of India [EMR/2015/001966] for extramural funding. SR acknowledges Department of Biotechnology, Govt. of India [BT/PR19201/BRB/10/1532/2016] for extramural funding. NB acknowledges the research fellowship received from CSIR, Govt. of India.

