## Supplementary material for "NMR-based analysis of nucleotide π-stacking in a crowded environment: Implications for prebiotic reactions": NMR_NT stacking_Supplementary File

### <u>Supplementary Information</u>

**Table S1. Experimental parameters, Δ and δ, that are used to record DOSY data for nucleotides at different concentrations**

| Concentration of Nucleotide (in mM) | Without PEG | | With PEG 8000 | |
|---|---|---|---|---|
| | Δ (ms) | δ (ms) | Δ (ms) | δ (ms) |
| **5'-AMP** | | | | |
| 10 | 60 | 2.5 | 60 | 4.5 |
| 50 | 60 | 2.6 | 60 | 4.5 |
| 100 | 60 | 2.8 | 60 | 4.8 |
| **5'-GMP** | | | | |
| 10 | 60 | 2.5 | 60 | 4.4 |
| 50 | 60 | 2.6 | 60 | 4.4 |
| 100 | 60 | 2.6 | 60 | 4.4 |
| **5'-CMP** | | | | |
| 10 | 60 | 2.5 | 60 | 3.8 |
| 50 | 60 | 2.6 | 60 | 4.0 |
| 100 | 60 | 2.8 | 60 | 4.0 |
| **5'-UMP** | | | | |
| 10 | 60 | 2.5 | 60 | 3.4 |
| 50 | 60 | 2.6 | 65 | 3.8 |
| 100 | 60 | 2.7 | 90 | 3.9 |

**Table S2. Diffusion constants for the nucleotides in the absence and presence of PEG**

| Concentration of Nucleotide (in mM) | Diffusion constant ($1*E^{-10}$) ($m^2/s$) without PEG | Diffusion constant ($1*E^{-10}$) ($m^2/s$) with PEG |
|---|---|---|
| **5'-AMP** | | |
| 10 | $5.08E^{-10} \pm 0.77E^{-12}$ | $1.75E^{-10} \pm 1.18E^{-12}$ |
| 50 | $4.84E^{-10} \pm 1.31E^{-12}$ | $1.62E^{-10} \pm 0.68E^{-12}$ |
| 100 | $4.51E^{-10} \pm 2.23E^{-12}$ | $1.64E^{-10} \pm 0.43E^{-12}$ |
| **5'-GMP** | | |
| 10 | $4.98\ E^{-10} \pm 2.18E^{-12}$ | $1.93E^{-10} \pm 2.20E^{-12}$ |
| 50 | $4.57E^{-10} \pm 2.42E^{-12}$ | $1.60E^{-10} \pm 0.94E^{-12}$ |
| 100 | $4.29E^{-10} \pm 0.60E^{-12}$ | $1.44E^{-10} \pm 0.45E^{-12}$ |
| **5'-CMP** | | |
| 10 | $5.53E^{-10} \pm 1.84E^{-12}$ | $1.44E^{-10} \pm 5.05E^{-12}$ |
| 50 | $5.29E^{-10} \pm 0.62E^{-12}$ | $1.98E^{-10} \pm 1.02E^{-12}$ |
| 100 | $5.06E^{-10} \pm 1.45E^{-12}$ | $1.96E^{-10} \pm 0.40E^{-12}$ |
| **5'-UMP** | | |
| 10 | $5.57E^{-10} \pm 4.22E^{-12}$ | $1.87E^{-10} \pm 5.03E^{-12}$ |
| 50 | $5.38E^{-10} \pm 1.59E^{-12}$ | $1.99E^{-10} \pm 2.15E^{-12}$ |
| 100 | $5.03E^{-10} \pm 1.29E^{-12}$ | $2.02E^{-10} \pm 0.19E^{-12}$ |

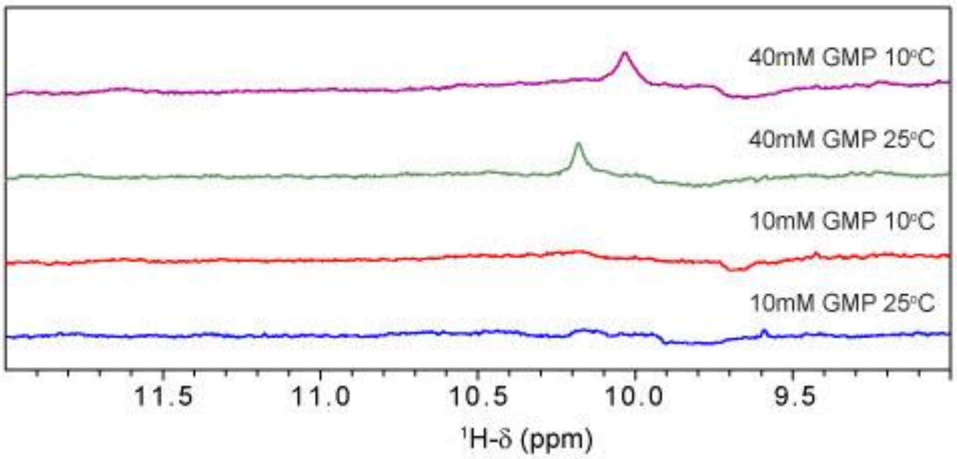


**Figure S1: Presence of G-quadruplex structure in 40 mM 5'-GMP concentration as indicated by the peak near 10 ppm in $^1H$ NMR**
